# Evidence for transient, uncoupled power and functional connectivity dynamics

**DOI:** 10.1101/2024.08.31.610630

**Authors:** Rukuang Huang, Chetan Gohil, Mark Woolrich

## Abstract

There is growing interest in studying the temporal structure in brain network activity, in particular, dynamic functional connectivity (FC), which has been linked in several studies with cognition, demographics and disease states. The sliding window approach is one of the most common approaches to compute dynamic FC. However it cannot detect cognitively relevant and transient temporal changes at the time scales of fast cognition, i.e. on the order 100 milliseconds, which can be identified with model-based methods such as HMM (Hidden Markov Model) and DyNeMo (Dynamic Network Modes) using electrophysiology. These new methods provide time-varying estimates of the “power” (i.e. variance) and of the functional connectivity of the brain activity, under the assumption that they share the same dynamics. But there is no principled basis for this assumption. In this work, we propose Multi-dynamic Network Modes (M-DyNeMo), an extension to DyNeMo, that allows for the possibility that the power and the FC networks have different dynamics. Using this new method on magnetoencephalography (MEG) data, we show intriguingly that the dynamics of the power and the FC networks are uncoupled. Using a (visual) task MEG dataset, we also show that the power and FC network dynamics are modulated by the task, such that the coupling in their dynamics changes significantly during task. This new method reveals novel insights into evoked network responses and ongoing activity that previous methods fail to capture, challenging the assumption that power and FC share the same dynamics.

**Highlights:**

- We show that our proposed model - Multi-dynamic Network Modes (M-DyNeMo) - infers transient, uncoupled dynamics for time-varying variance (i.e. power) and functional connectivity (FC) in MEG data.
- M-DyNeMo infers plausible power and FC networks that are reproducible across different subjects, datasets, and parcellations.
- M-DyNeMo reveals task modulates the dynamics of the power and the FC networks, which induces a coupling between dynamics.
- The evoked FC network response to a (visual) task is different to the evoked network response in power.
- The datasets and scripts for performing all the analysis in this paper are made publicly available.

## 1 Introduction

Recently, researchers have adopted the view that cognitive tasks are performed not by individual brain regions working in isolation, but by networks of several brain areas which communicate with each other. A metric often used to characterize this communication is functional connectivity (FC), which is defined as the temporal correlation of functional activity between spatially remote regions ([Friston, 1994]). The study of FC has become increasingly popular as evidence suggests that FC is related to underlying neural activity ([Nir et al., 2008]) and significant differences have been observed in FC between healthy and diseased states ([Greicius, 2008, Menon, 2011, Heine et al., 2012]).

Traditionally, it was typically assumed that the strength of interactions between regions is constant over time (stationary) during resting-state scans. However, given the brain’s dynamic nature ([Rabinovich et al., 2012]), an increasingly important perspective is how functional networks change over time. One of the most common approaches to compute dynamic FC is the sliding window approach ([Chang and Glover, 2010, Allen et al., 2014, Liuzzi et al., 2019]). Clustering methods like K-means clustering are followed to identify reproducible and transient patterns of FC states ([Allen et al., 2014]).

Despite its popularity, the sliding window approach is a heuristic approach in which the choice of hyperparameters can have an impact on the interpretation of results. More importantly, sliding window methods are deficient at adapting to fast dynamics in the data and finding sub-second transitions in brain dynamics, which have been shown to be present in electrophysiological data like magnetoencephalography (MEG) ([Vidaurre et al., 2016, Vidaurre et al., 2018]). This is due to fixed window size being assumed, with larger windows being insensitive to fast changes, and smaller windows giving noisier estimates of FC.

Recent advances in the field of machine learning provide methods for inferring dynamic FCs in a data-driven way. An example is the recently proposed Dynamic Network Modes model (DyNeMo, [Gohil et al., 2022]) which allows for co-activation of multiple networks simultaneously. It has been shown that DyNeMo can infer dynamic FC more accurately than state-based models, such as the Hidden Markov Model ([Gohil et al., 2022]), and can identify more parsimonious dynamic network descriptions when studying task data ([Gohil et al., 2024b]).

One important assumption that underlies DyNeMo is that time-varying (TV) variance (which we will refer to equivalently as “power”) and FC follow the same temporal dynamics, but there is no principled reason to assume fluctuation in connections between brain regions should be coupled with that in power of individual brain regions. This motivates us to present Multi-dynamic Network Modes (M-DyNeMo), an extension to DyNeMo, to infer potentially uncoupled power and FC dynamics.

In this paper, with resting-state MEG data, we present evidence of separate power and FC dynamics given by the traditional sliding window approach, and show that plausible networks of separate dynamics can be inferred by M-DyNeMo. We further demonstrate that the networks are reproducible across different subjects, datasets, and parcellations, validating the robustness of M-DyNeMo on MEG data. Using visual task data, M-DyNeMo reveals distinct evoked network responses of power and FC dynamics to visual stimuli. We also show evidence for task-induced coupling between power and FC dynamics.

## 2 Results

In this section, we study three real MEG datasets (Section 6.1): MEGUK-38 (resting-state), MEGUK-52 and Wakeman-Henson.

### 2.1 Sliding window approach reveals distinct power and FC dynamics

Firstly, we apply the well-established sliding window approach followed by K-Means clustering to identify two separate dynamics according to TV power (variance) and TV FC (correlation) respectively (See Section 6.3, [Allen et al., 2014]) from amplitude time courses (1-45Hz) of 38 brain regions obtained from MEGUK-38 resting-state data. Figure 2.1a illustrates the first 20 seconds of the state time courses (K-Means cluster assignments) for the first subject in the MEGUK-38 resting-state data. Qualitatively, we can see that TV power and FC does not follow the same dynamics, and the correlation between the two time courses is low, as shown in Figure 2.1b. Furthermore, each of the two state time courses corresponds to reasonable power and FC maps respectively, demonstrated in Figure 2.1c.

**Figure 2.1:**
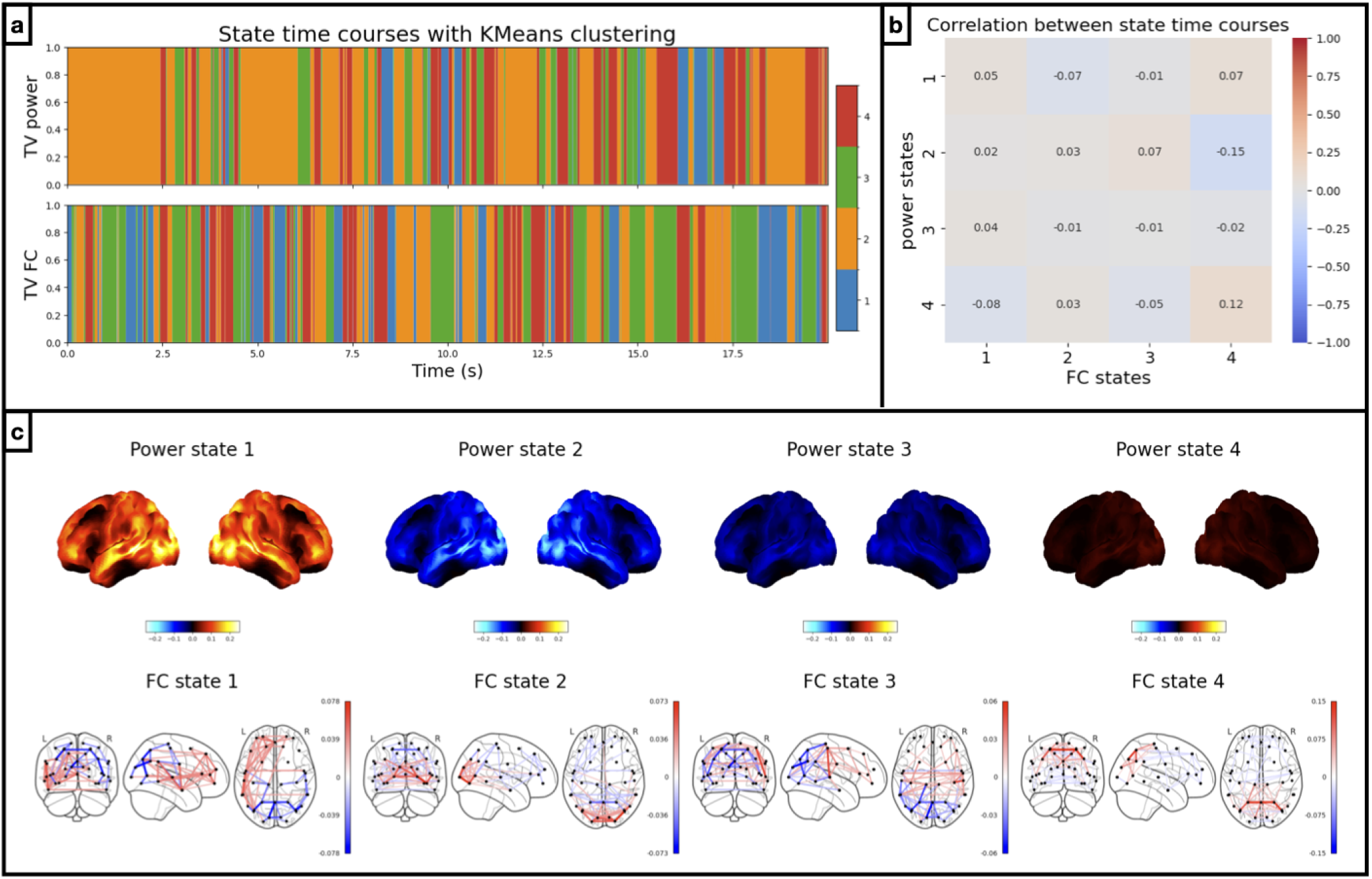
Sliding window approach reveals distinct power and FC dynamics. *Results from amplitude time courses (1-45Hz) of 38 brain regions obtained from the MEGUK-38 resting-state data*. a) K-Means state time courses for TV variance (i.e. power) (top row) and correlation (i.e. FC) (bottom row), using 2s sliding windows. Different colours indicate activation of different brain states and only the first 20s of the first subject is shown. b) Correlation between power (y-axis) and FC state time courses (x-axis). c) Power maps of each K-Means state are plotted on the top row. Red areas indicate brain areas with higher power (and blue areas lower power) than the average across states. FC maps are plotted on the bottom row. Red edges indicate positive correlations (and blue edges negative correlations) between brain areas.

**Figure 2.2:**
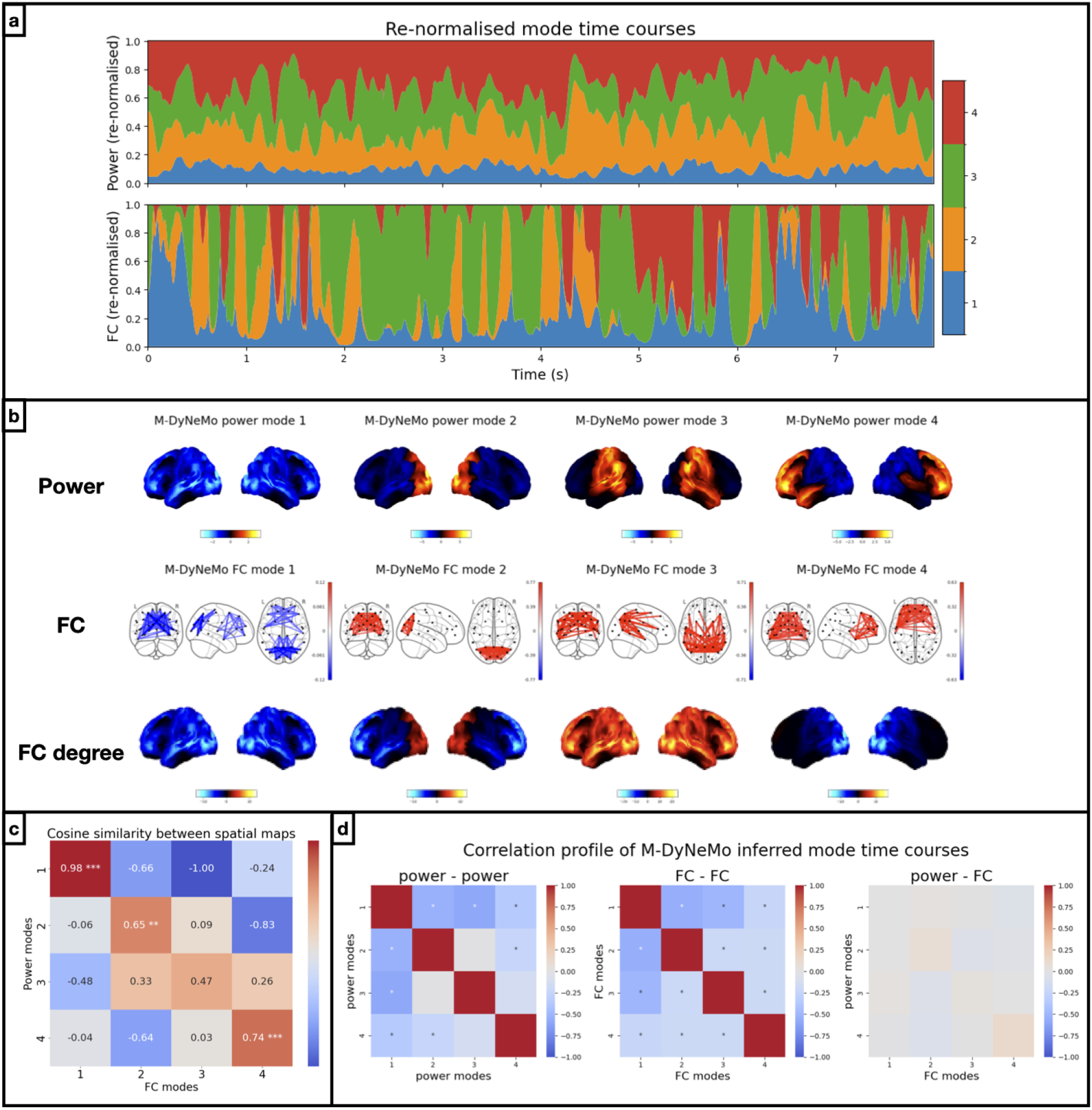
Power and FC have similar spatial patterns but distinct dynamics. *Results of running M-DyNeMo on the amplitude time courses (1-45Hz) of 38 brain regions obtained from the MEGUK-38 resting-state data*. a) Inferred mode time courses (renormalised) for power, *α* (top) and FC, *β* (bottom). b) Top row shows power spatial modes where red areas show higher (blue areas show lower) activation than average across modes. Middle row shows FC spatial modes where red edges show positive (and blue edges show negative) connectivity. Bottom row shows the FC degree maps where red areas show higher (blue areas show lower) node degree (i.e. centrality) than average across modes. c) Pairwise cosine similarity between power and FC degree spatial maps. Diagonal entries with significantly large values are annotated with asterisks depending on the p-values. d) Correlation profile of mode time courses for within power modes (left), within FC modes (middle) and between power and FC modes. Correlations significantly (at 5% significance level) differ from zero under a maximum statistic permutation test are marked with an asterisk.

While Figure 2.1 provides preliminary evidence that power and FC have distinct dynamics, we do not know if this is an artefact of the choice of arbitrary hyperparameters (e.g. window size) or the inability of sliding window approaches to automatically adapt over time to fast dynamics (Section A.2 demonstrates this on simulated data). We instead need a method that can infer multiple dynamics in an adaptive, data-driven way. This motivates a new approach, Multi-dynamic Network Modes (M-DyNeMo).

### 2.2 Power and FC have similar spatial patterns but distinct dynamics

A formal description of the M-DyNeMo is available in the Methods section (Section 6), including the generative model and the inference framework to fit the model to data. Conceptually, M-DyNeMo learns two basis sets of spatial configurations of networks (called *modes*), one for power (where the network is a spatial map over brain regions) and another for the FC (where the network is an edge-wise connectome). The underlying assumption is that the TV power/FC is a time-varying linear mixture of the basis set. The time series of mixing coefficients characterises the dynamics of the modes and is referred to as the *mode time courses*. Fluctuations in power and FC can be described independently via dynamics in the mixing of their respective basis sets. We can show using simulated data that M-DyNeMo performs more accurate inference than standard DyNeMo, if the ground truth contains distinct dynamics for power and FC (see Section A.2).

First, we trained M-DyNeMo on the amplitude time courses (1-45Hz) of 38 brain regions obtained from MEGUK-38 resting-state data. Figure 2.2a shows the re-normalised mode time courses for the first 8 seconds of the first subject in this dataset (See Section E.1 for details on re-normalisation). We see qualitatively that mode time courses for power (*α*) and for FC (*β*) possess very different characteristics, in the sense that *β* is much more binarised than *α*.

Power and FC networks for each mode are shown in Figure 2.2b. These networks resembled those found in multiple previous studies ([Baker et al., 2014, Vidaurre et al., 2016, Gohil et al., 2022, Gohil et al., 2024b]). For a clearer visual comparison, we also show the FC degree maps, each of which plot the sum of correlations for each node, relative to the average across modes. Note, the ordering of spatial modes found by the model is arbitrary, hence the network maps shown in this figure have been re-ordered post-hoc to match the spatial patterns of activity in power and FC for each pair of modes. Even though there is no requirement for them to be similar, the power and the FC spatial maps have highly corresponding spatial patterns. Specifically, there is significantly high cosine similarities between power and FC degree maps for mode pairs 1, 2 and 4 under a maximum statistic permutation test (see Section E.2 for details), illustrated by the diagonal entries in Figure 2.2c.

Although we identify power and FC modes that show activity in the same regions, their corresponding mode time courses (dynamics, *α* vs *β*) are not significantly correlated (under a maximum statistic permutation test across subjects), as demonstrated in Figure 2.2d, plotted on the right, suggesting time points with high power in a particular region may not necessarily also have high FC in the same region. The correlations within the power mode dynamics (*α*, left) and within the FC mode dynamics (*β*, middle) are mostly negative, indicating when a mode activates the others deactivate.

In summary, Figure 2.2 shows that power and FC have similar spatial patterns but distinct dynamics. However, one concern is that the distinct dynamics might be enforced by the assumptions made by M-DyNeMo. To mitigate this concern, we used simulations in Section A.3 to show that if the data is generated using a single, shared dynamic for power and FC, then M-DyNeMo does not incorrectly infer distinct dynamics.

### 2.3 Single dynamic DyNeMo is dominated by time-varying power

Given the evidence that there are distinct dynamics, we were interested to learn which of the dynamics dominate the decomposition when only a single dynamic is allowed, as is the case in standard DyNeMo. To investigate this we assumed that the power and FC share a single dynamic given by the power mode time course *α*, extracted from a trained M-DyNeMo, and then re-calculated the power and FC for each mode. The resulting networks are shown in Figure 2.3a. Comparing this with the networks inferred by standard DyNeMo on the same amplitude time courses (1-45Hz) of 38 brain regions from the MEGUK-38 resting-state data, shown in Figure 2.3b, we can see that the power mode time course (*α*) inferred by M-DyNeMo contains similar information as the mode time course inferred by standard DyNeMo. Quantitatively, for each pair of modes, there is a significantly large cosine similarity (under a maximum statistic permutation test) between the re-calculated networks using M-DyNeMo inferred power mode time course *α* and the inferred networks given by standard DyNeMo. This suggests the description provided by standard DyNeMo is dominated by the dynamics in power rather than FC. Furthermore, by re-calculating networks using M-DyNeMo inferred FC mode time course *β*, we show in Figure B.1 that the FC dynamics given by M-DyNeMo contains different information as the mode time course inferred by standard DyNeMo.

**Figure 2.3:**
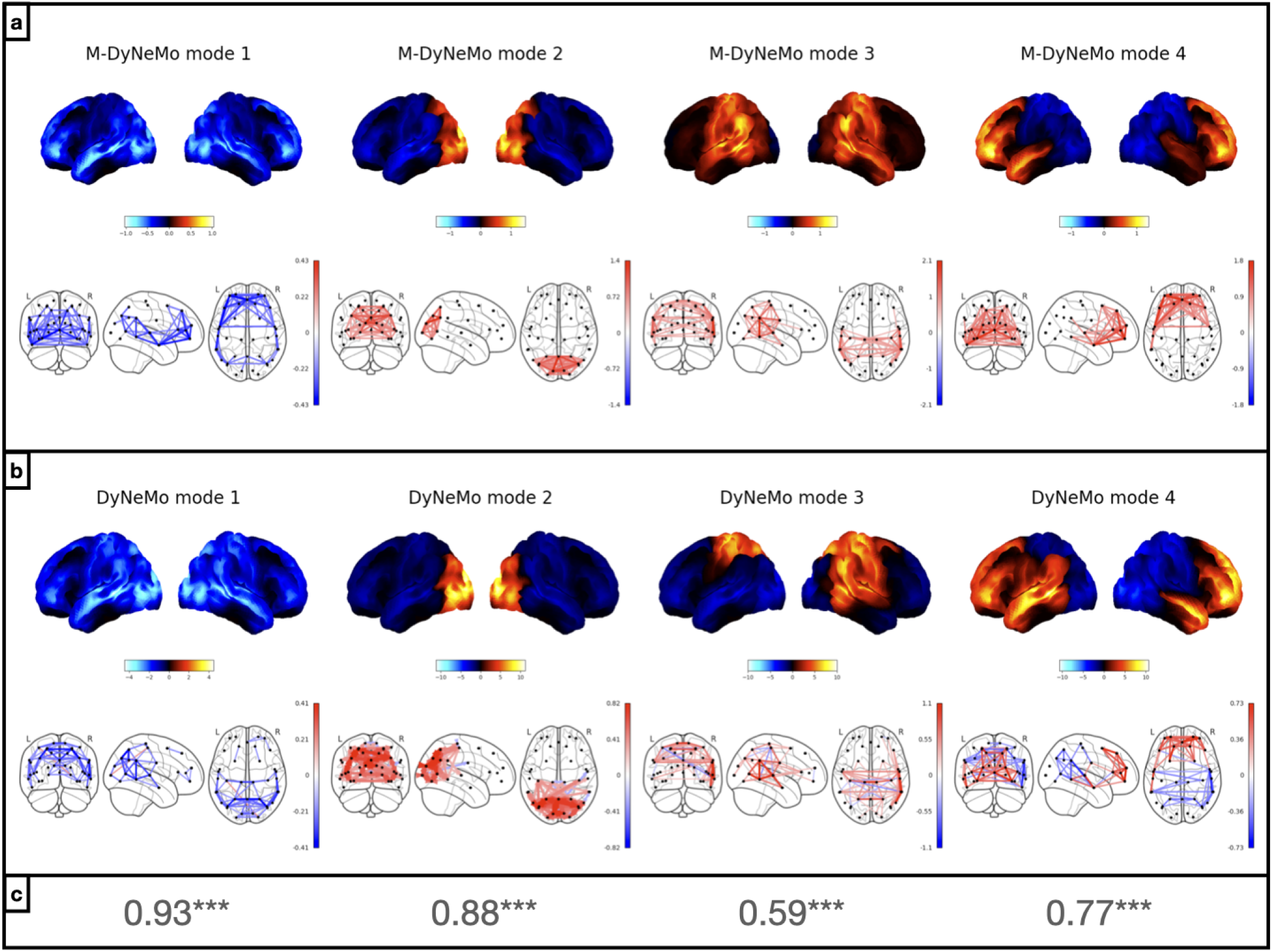
Single dynamic DyNeMo is dominated by time-varying power. *Results from amplitude time courses (1-45Hz) of 38 brain regions obtained from MEGUK-38 resting-state data*. a) Power (top row) and FC (covariance, bottom row) network maps given by re-calculating time varying covariances on M-DyNeMo inferred power mode time course *α*. b) Power (top row) and FC (covariance, bottom row) maps given by DyNeMo. c) Cosine similarity between re-calculated covariances on M-DyNeMo inferred power mode time course *α* and DyNeMo inferred covariances for each pair of modes.

### 2.4 M-DyNeMo inferred networks are reproducible across subsets of data

To evaluate the reproducibility and robustness of the model, we train M-DyNeMo on different subsets of subjects of the MEGUK-38 resting-state dataset. We split 65 subjects of this dataset into two halves where the first half contains subjects 1 to 32 and the second half contains subjects 33 to 65. M-DyNeMo is trained independently on each half. Figure 2.4a and b shows that network maps inferred by M-DyNeMo on each half, with modes matched using cosine similarity, which is also shown in the figure for each pair of modes. In general, we see the same networks in each half with significantly large cosine similarity under a maximum statistic permutation test. There are only slight differences in the spatial pattern of activity in power modes 2 and 3.

**Figure 2.4:**
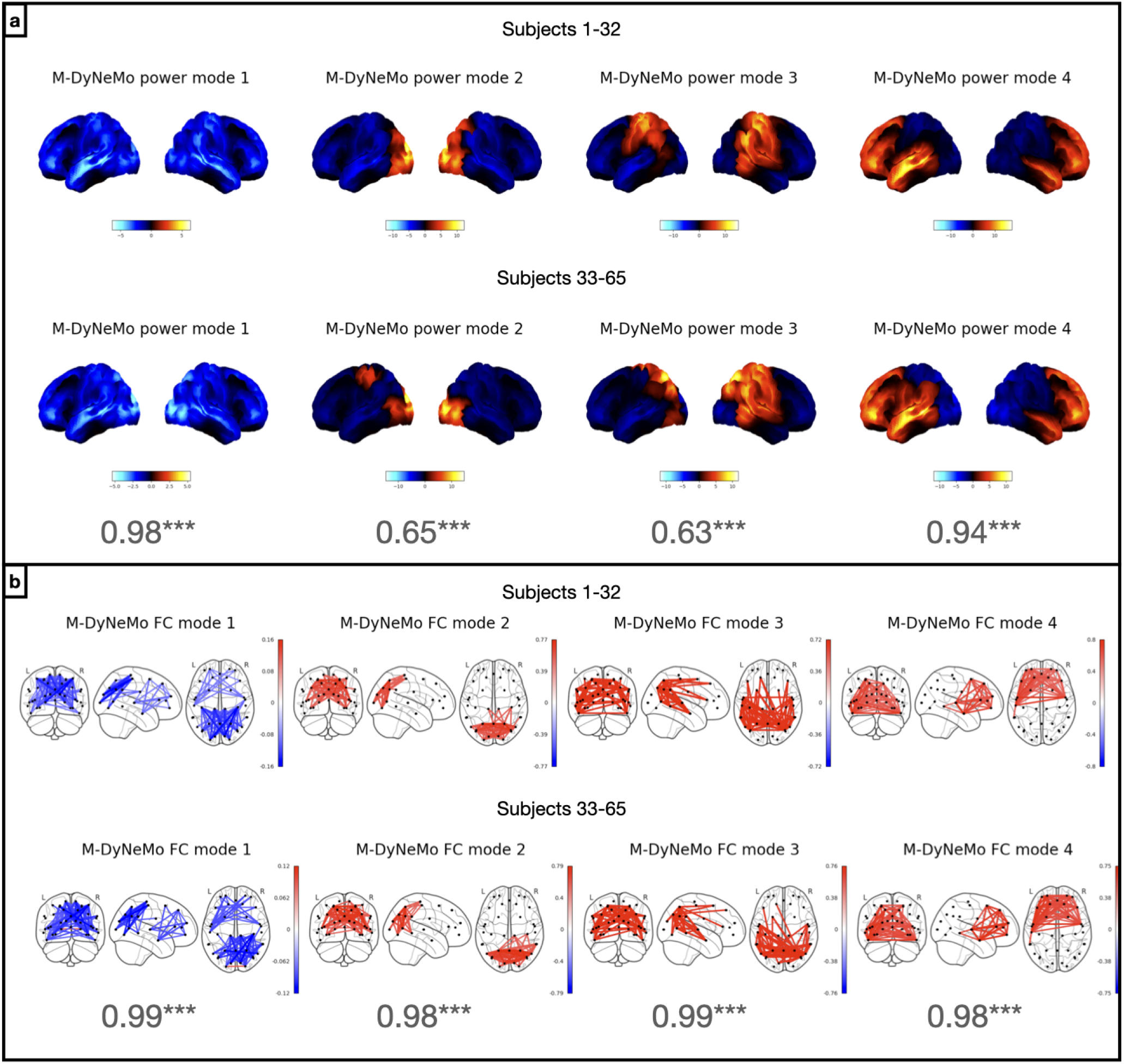
M-DyNeMo inferred networks are reproducible across halves. *Results from amplitude time courses (1-45Hz) of 38 brain regions obtained from MEGUK-38 resting-state data*. a) Power spatial modes given by training M-DyNeMo on subjects 1-32 (top row) and subjects 33-65 (bottom row) independently. The cosine similarity between power maps from each half is also shown for each pair of modes. b) FC spatial modes given by training M-DyNeMo on subjects 1-32 (top row) and subjects 33-65 (bottom row) independently. The ordering of both power and FC modes are matched with cosine similarities between networks from each half, which are shown in the figure for both power and FC networks.

### 2.5 Power and FC dynamics show different timings in evoked network response

In Section 2.2, we saw M-DyNeMo infers distinct dynamics (i.e. uncorrelated mode time courses) for power and FC using the amplitude time courses (1-45Hz) from resting-state MEG data. We now turn to task MEG data and ask the question: Do *α* and *β* respond differently to tasks? We perform this investigation with the Wakeman-Henson dataset, where participants are presented with images of famous, unfamiliar, or scrambled faces. This allows us to study how the mode time courses respond to visual stimuli as well as the difference in responses to different visual stimuli.

The network plots are shown in Figure 2.5a, which are comparable to those inferred from MEGUK-38 resting-state data, shown in Figure 2.2b. In Figure 2.5b, the evoked network responses are shown for different contrasts, including the average of all visual stimuli, the difference between faces (including both famous and unfamiliar faces) and scrambled faces, and the difference between famous and unfamiliar faces.

**Figure 2.5:**
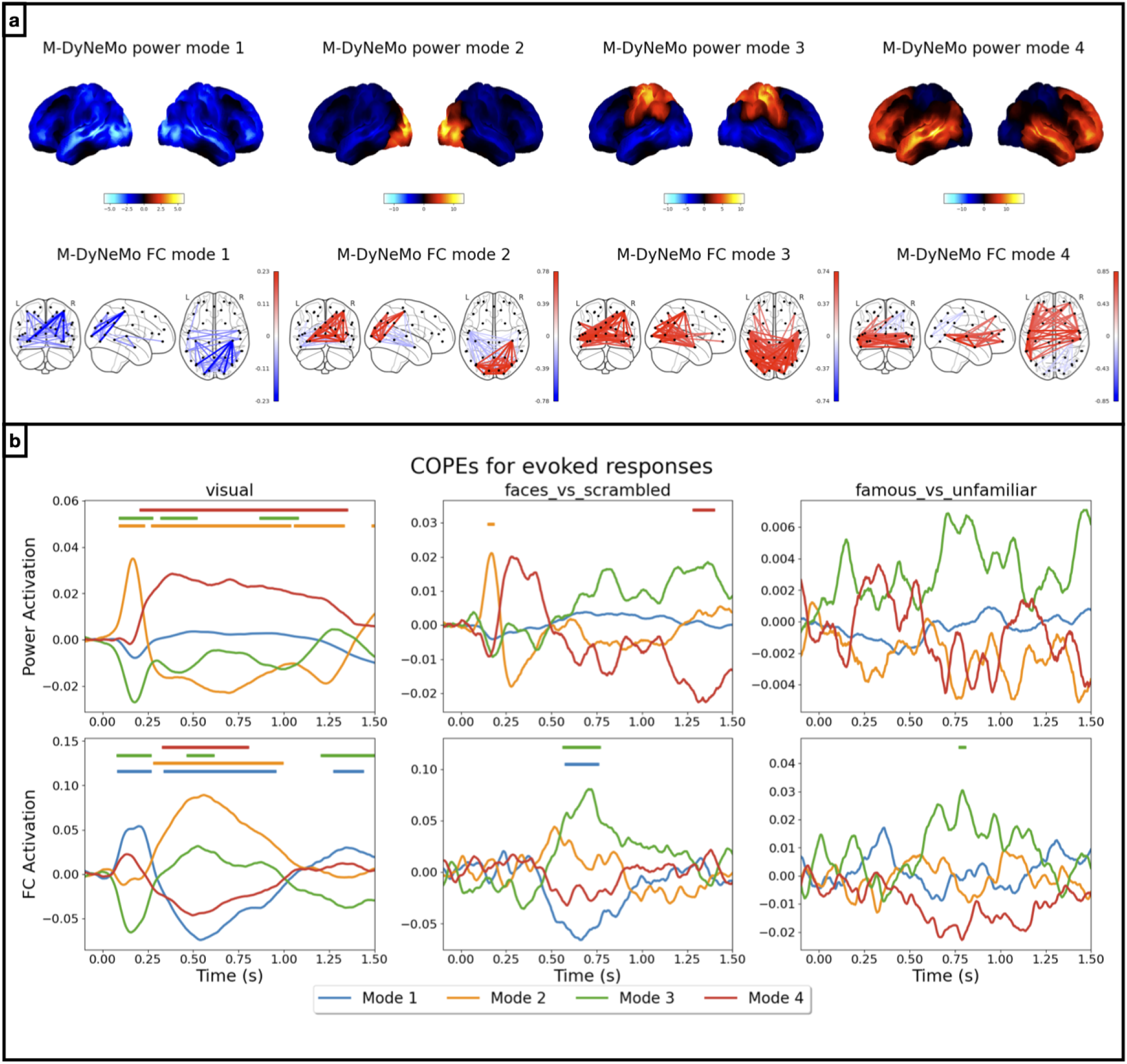
Evoked responses have different timings in different mode time courses. *Results from amplitude time courses (1-45Hz) of 38 brain regions obtained from Wakeman-Henson data*. a) Power spatial modes are shown on the top row and FC spatial modes are shown on the bottom row. b) Evoked responses for re-normalised *α* (top row) and *β* (bottom row) for different contrasts on different columns, including the average visual response (left), the difference between faces and scrambled faces (middle), and the difference between famous and unfamiliar faces (right). Here different colours represent baseline corrected evoked responses for different modes as contribution to either the overall variance (for *α*) or the overall correlation (for *β*). Periods with significant (de-)activation are indicated with a bar for each mode.

#### 2.5.1 Power dynamics

We focus on the top row of Figure 2.5b first. Similar to the analysis performed in [Gohil et al., 2024b], a two-level General Linear Model (GLM, [Winkler et al., 2014]) is used for computing evoked network responses, where an evoked response analysis is carried out on each of the task epoched M-DyNeMo mode time courses. Note, that this is carried out *after* M-DyNeMo is trained, i.e. M-DyNeMo has no knowledge of the task timings. A maximum statistic permutation test is used to identify periods of significant responses.

We see in the top row of Figure 2.5b that evoked network responses for power, *α*, are broadly consistent with those given by training standard DyNeMo on the same data (see Figure B.3). For example, in the average visual response, there is an immediate activation of the visual network (*α* mode 2) followed by an activation of the frontal network (*α* mode 4) and a deactivation of the visual network. In the difference between unscrambled faces (famous plus unfamiliar) and scrambled faces, there is an activation of the visual network and a delayed deactivation of the frontotemporal network. The only difference is that there is no activation of the suppressed network (*α* mode 1) at around 600ms after the onset like in the analysis of standard DyNeMo. There are no significant differences when we look at the responses to famous and unfamiliar faces.

#### 2.5.2 FC dynamics

We now draw our attention to the bottom row of Figure 2.5b where the evoked responses of the FC network mode dynamics, *β*, are illustrated. For the average visual response (left), we see an immediate activation of mode 1 and deactivation of mode 3, followed by a delayed and persistent activation of mode 2 as well as a deactivation of the mode 1 and mode 4. Looking at the differences in response to faces and scrambled faces, there is only a delayed activation of mode 3 and a deactivation of the mode 1. Finally, there is a short-lived activation of the mode 3 when comparing responses to famous and unfamiliar faces.

### 2.6 The coupling between power and FC dynamics is modulated by task

As shown in Figure 2.5, there is evidence of different timings in evoked network responses for power and FC dynamics. This motivated us to investigate whether the extent to which there is coupling, between power dynamics and FC dynamics, is different between task and rest. To this end, we train M-DyNeMo on a new dataset, MEGUK-52, which contains 63 subjects that have both resting-state and visual task recordings, totalling 126 sessions. Figure 2.6a shows the inferred networks, which are broadly consistent with those inferred on the same dataset, using a different parcellation compared to in Section 2.2 (MEGUK-38 resting-state data) and on a different dataset to Section 2.5 (Wakeman-Henson - which only contains visual task data and on a different set of subjects). This again demonstrates the reproducibility of M-DyNeMo on different datasets and parcellations.

**Figure 2.5:**
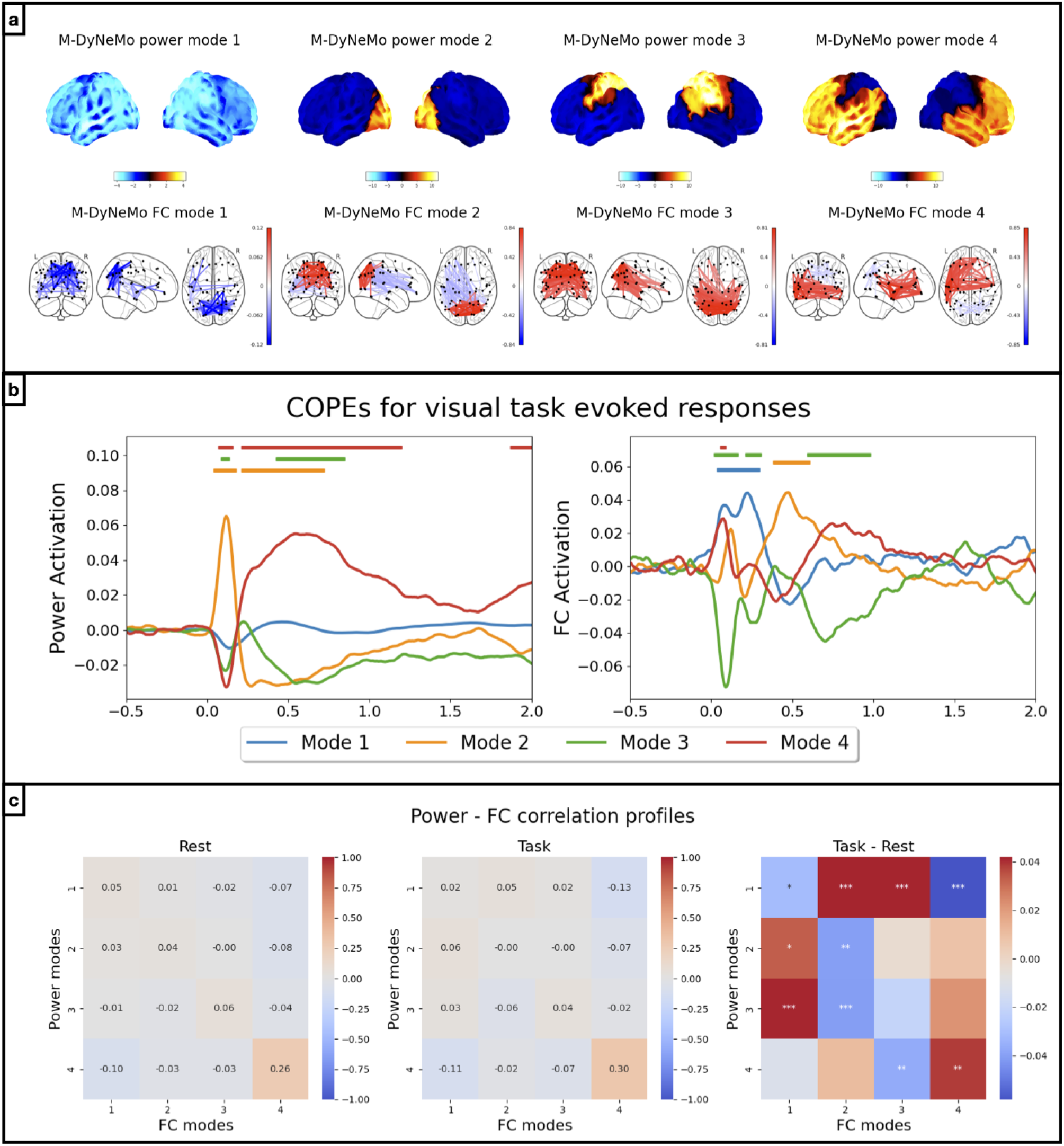
Coupling between *α* and *β* is modulated by task stimuli. *Results from amplitude time courses (1-45Hz) of 52 brain regions obtained from MEGUK-52 data*. a) Power (top row) and FC (bottom row) spatial modes are plotted. b) Evoked responses (baseline corrected) for re-normalised *α* (left) and *β* (right) to visual stimuli. Different colours show evoked responses for different modes. c) Correlations between *α* and *β* during rest (left) and during task (middle) are shown. The right plot illustrates difference between the correlation profiles where significant differences are marked with asterisks.

The evoked network responses to visual stimuli, shown in Figure 2.6b, which is consistent with the findings in Section 2.5. Figure 2.6c shows the correlations between the power (*α*) and FC (*β*) dynamics during rest (left), those during epochs of 0.5 seconds before and 2 seconds after the visual stimuli (middle), as well as the difference in correlations between task and resting state. In particular, from rest to task, we see increased correlation of *α* mode 2 with *β* mode 1 and decreased correlation with *β* mode 2, which is consistent with the immediate activation of *α* mode 2 and *β* mode 1, and the delayed activation of *β* mode 2 in Figure 2.6b. It is worth noting that the sliding window approach on this dataset also provides evidence for the difference in the coupling of the two dynamics (See Figure B.4).

## 3 Discussion

With resting-state MEG data, we found preliminary evidence for the existence of distinct power and FC dynamics using a standard sliding window approach. However, this approach lacks the ability to adapt to transient dynamics at unknown time-scales in electrophysiological data. We demonstrated the deficiency of sliding window approaches to do this using simulated data in Section A.2. Therefore, we proposed a new generative model approach that automatically tunes to the required time-scales based on the data, and that can incorporate multiple dynamics: M-DyNeMo.

Using M-DyNeMo, we show that while they have similar spatial patterns, the two mode time courses, one for power and one for FC, possess very different characteristics and there is no significant correlation between them, i.e. they are decoupled. The same property can also be found in two other datasets (see Figure B.5). It should be noted that there is no constraint imposed by the generative model or inference framework in M-DyNeMo on the dynamics. It is solely the data itself that decides the dynamics are decoupled and the FC dynamics is more binarised than the power dynamics.

Using standard single-dynamic DyNeMo, consistent differences have previously been found in mode time courses between young and old participants ([Gohil et al., 2024a]). Therefore, a future direction is to see if the FC mode time courses, which to date will have been ignored due to its relative low signal in real data, can provide more understanding into aging, and distinctions between healthy and diseased brains.

Aside from resting-state data, we also explore the additional insights M-DyNeMo provides on task data. In Section 2.5, we show that the two mode time courses respond differently to visual stimuli. Moreover, we provide evidence that coupling between power and FC dynamics is modulated by stimuli. This could be related to the findings in [Barttfeld et al., 2015] that complexity in FC dynamics is inversely proportional to the level of consciousness. Overall, our results demonstrate how M-DyNeMo has opened a new door to studying the relationship between cognition and dynamic FC.

In Section 2.3, we showed that standard, single-dynamic DyNeMo tends to ignore the FC dynamics in real data, and is dominated by the power dynamics. This is consistent with the idea that the signal from time-varying FC is weaker in real data than that from power, which is perhaps to be expected given that power is easier to estimate in general. If we look at the fluctuation of the reconstructed time-varying covariances from both M-DyNeMo and standard DyNeMo (Figure B.2), we can see that the shared-dynamics constraint of standard DyNeMo causes lower variability in FC, which is consistent to the findings of applying multi-dynamic modelling to fMRI data ([Pervaiz et al., 2022]).

In Section A.3, we showed that if the underlying data is generated by a single dynamic process, then M-DyNeMo does not enforce distinct dynamics for FC and power, and correctly infers a single, shared set of dynamics. It is worth noting that although data are simulated using the generative model of a Hidden Markov Model, where time courses are binarised, M-DyNeMo is still capable to learn from the data and infer binarised time courses, without a constraint being posed on its generative model. We have also explored using an edge time series technique proposed by [Faskowitz et al., 2020] on the simulated dataset. However, we find it gives noisy estimates of FC and fails to capture the transient temporal changes accurately.

In this paper, we used amplitude envelope time courses (1-45Hz) from MEG data, and correspondingly used correlation between amplitude envelopes as a measure of FC. However, this ignores more detailed spectral information in the MEG signal. For example, the phase-locking between brain regions (coherence) has recently gained increasing interest as a measure of FC in electrophysiology data. [Vidaurre et al., 2016] introduced a method called *time-delayed embedding* for including spectral information into the covariance matrix of the time-delay embedded data. However, it is not trivial to decompose this covariance matrix into parts that correspond to power and parts that correspond to FC, similar to what has been done in Section 6.4. The additional PCA step for reducing dimensionality complicates the modelling even further. An important aspect of future work is to find a way to allow inference of separate dynamics based on spectral information.

## 4 Conclusions

We proposed a new method for inferring potentially separate power and FC dynamics. We show that the proposed model is robust and inferred networks are reproducible across subjects, datasets and parcellations. We use this to show that power and FC have similar spatial patterns, but distinct dynamics, in ongoing MEG brain activity. Furthermore, we show that the two types of dynamics respond differently to visual stimuli in MEG task data, and that the extent to which there is coupling between the two types of dynamics is modulated by task.

## 5 Data and code availability statement

Data used are publicly available. Availability of MEGUK is at the official site https://meguk.ac.uk/database/ and for Wakeman-Henson, we refer the readers to the original paper [Wakeman and Henson, 2015]. Source code for M-DyNeMo is available in the *osl-dynamics* toolbox ([Gohil et al., 2024a]) and scripts to reproduce results in this manuscript are available at:

https://github.com/OHBA-analysis/Huang2024_EvidenceForUncoupledDynamics.

## 6 Methods

### 6.1 Dataset

Two real MEG datasets are used in this paper. The first dataset is part of the UK MEG Partnership acquired using a 275-channel CTF MEG scanner at a sampling frequency of 1.2kHz. This dataset includes (eyes open) resting-state recordings of 65 subjects and visual task recordings of 67 subjects, of which 63 subjects have both resting-state and task MEG recordings. We refer to this dataset as the **MEGUK** dataset.

The second dataset is a visual task MEG dataset collected using an Elekta Neuromag Vectorview 306 scanner at a sampling frequency of 1kHz. In this dataset, 19 participants (11 male, 8 female) were scanned 6 times, during which they were presented with an image of famous, unfamiliar, or scrambled faces. The MaxFiltered data is publicly available and we direct the readers to the original paper ([Wakeman and Henson, 2015]) for more details on the experimental design and data collection. We refer to this dataset as the **Wakeman-Henson** dataset.

#### Data preprocessing

Both datasets are preprocessed using the OHBA software library (OSL, [Quinn et al., 2022]) with the same pipeline. A band-pass filter from 0.5Hz to 125Hz and a notch filter with 50Hz and 100Hz were applied to the raw data. Then the data was downsampled to 250Hz, after which automatic bad segment and bad channel detection were used to remove abnormally noisy segments and channels of the recordings. A final independent component analysis (ICA) step with 64 components was used to remove noise.

#### Source reconstruction

Coregistration and source reconstruction were done using OSL. MEG data were firstly coregistered with the structural MRI data and digitised headshape points acquired with a Polhemus pen of each subject. Then the data were source reconstructed onto an 8mm isotropic grid using a linearly constrained minimum variance (LMCV) beamformer ([Van Veen and Buckley, 1988, Van Veen et al., 1997]).

#### Parcellation

Two atlases are used for parcellating the MEGUK dataset, one is the Giles parcellation with 38 regions of interest (ROIs) and the other is the Glasser parcellation with 52 ROIs, referred to as **MEGUK-38** (in which only the resting-state recordings are used in this paper) and **MEGUK-52** respectively. The Wakeman-Henson dataset is parcellated with the Giles parcellation to 38 ROIs. Finally, the symmetric multivariate leakage reduction ([Colclough et al., 2015]) and sign-flipping algorithm ([Vidaurre et al., 2016]) were applied.

#### Preparation before model training

Extra steps are taken to help the convergence of model training. Data were band-pass filtered between 1 and 45Hz to concentrate the data in the frequency range where neural activity is the most prominent before the amplitude envelope signals were extracted using Hilbert transform ([Feldman, 2011]). Non-overlapping moving average of window size of 25 samples (100ms at 250Hz) is applied to smooth the amplitude envelope data. Then a standardisation step (z-transform) is applied to ensure zero mean and unit variance for each channel. This is done independently for each recording session. Finally, we orthorgonalise the data with a full rank PCA (See Section 6.4.1 for details).

### 6.2 Measure of power and FC

In this paper, variance, instead of mean activation, of the amplitude of the recording signals is used as a measure of power. This is because we adopt the so-called *zero-mean model* in Section 6.4, which has been widely used in previous literature ([Vidaurre et al., 2016, Vidaurre et al., 2018, Gohil et al., 2022]). Correlation between amplitude signals is used as a measure of FC.

### 6.3 Sliding window approach

The aim of this method is to use a model free approach to partition the time dimension into a finite number of states according to either dynamic changes in power or FC. The heuristic approach is a combination of sliding window and K-means clustering. The steps include:

- Apply sliding window with window size of 500 samples (2s at 250Hz) and step size of 10 samples (40ms at 250Hz).
- For each window, calculate the correlation matrix and the standard deviations of the data.
- Apply K-means clustering algorithm to the time-varying correlation matrices and standard deviations separately.

Each time point is assigned to a cluster, which in our case can be thought of as a state, according to either TV power or FC. Hence we have formed two state time courses for power and FC separately.

### 6.4 Generative model of M-DyNeMo

#### Notations

- [*n*] = {1, … *n*} for *n* ∈ ℕ.
- ***x*** is a column vector.
- ***A*** is a matrix.
- ***I***_*n*_ is an *n* × *n* identity matrix.
- *x*_1:*n*_ = {*x*_*i*_ : *i* ∈ [*n*]}.

In this section we formulate the generative model of M-DyNeMo. Similar to DyNeMo, it is assumed that the data is generated by a multivariate Gaussian distribution with zero mean. Let ***x***_1:*T*_ be the observed data, where *T* is the number of samples/timestamps and ***x***_*t*_ is a vector of length *N*_*c*_ - the number of channels. Then

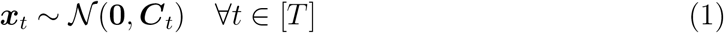

independently, where ***C***_*t*_ is the time-varying covariance matrix of the data at time *t*.

In M-DyNeMo, it is further assumed that ***C***_*t*_ can be decomposed into

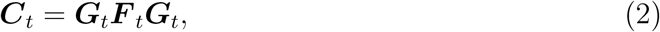

where ***G***_*t*_ is a diagonal matrix with strictly positive diagonal entries and ***F*** _*t*_ is a positive-definite matrix with ones on the diagonal. In particular, the diagonal elements of ***G***_*t*_ are the time-varying standard deviations and the off diagonal elements of ***F*** _*t*_ are the time-varying correlations of the data at time *t*. The time-varying standard deviations and correlations are assumed to be generated by a linear mixture of a finite number of modes:

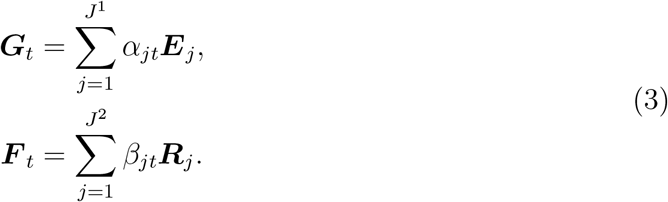

Here ***E***_*j*_ is a diagonal matrix with positive diagonal entries, and ***R***_*j*_ is a positive-definite matrix with ones on the diagonal. Notice here that the number of modes *J* ^1^, *J* ^2^ can be different. The mixing coefficients *α*_*jt*_ and *β*_*jt*_ are generated through a s of tmax transformation of latent probabilistic variables 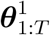, 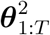, respectively, whose prior distribution is parameterised by an RNN:

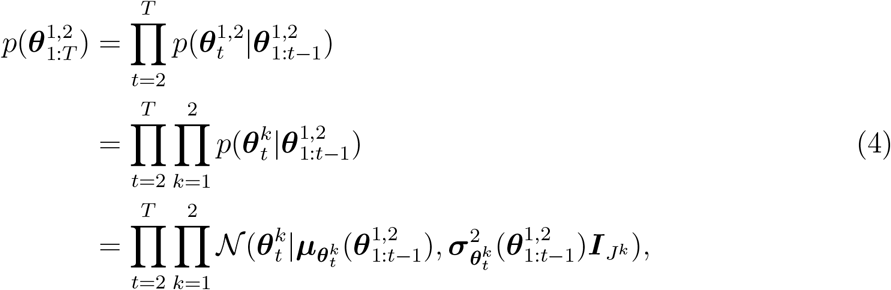

where

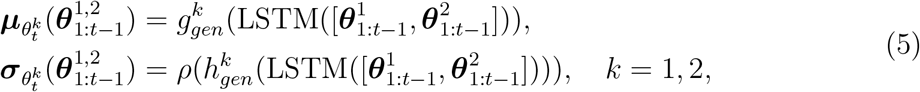

and 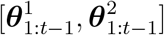 represents concatenating 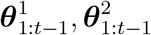 along the mode dimension. Here LSTM is a long short-term memory recurrently neural network ([Hochreiter and Schmidhuber, 1997]), 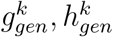 are learnable linear transformations (dense layers), and *ρ* is the softplus activation.

With this generative model, each of the latent logits 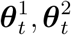 are generated by the history of both logits. This allows information to be communicated between the two different time courses in the generative model. An overview of the generative model is illustrated in Figure C.1.

#### 6.4.1 Orthogonalisation with PCA

Due to model complexity, the stability of training the generative model above is suboptimal, especially on small datasets. We found in practice that orthorgonalising the data, i.e. removing the correlation between channels, improved convergence of model training significantly. To this end, a full rank Principal Component Analysis (PCA) was applied to the data. Notice that the PCA-projected channels are linear combinations of the original channels, which makes separating dynamics of power and FC in this projected space non-trivial. Hence we instead modify the generative model and incorporate the PCA projection matrix into the generative model specified above. This allows easy interpretation of the learnt observation model parameters.

Formally, let 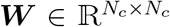 be the matrix with eigenvectors of the covariance matrix of the data on the rows, then the PCA-transformed data 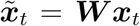 at time *t* has the following distribution, under the M-DyNeMo generative model:

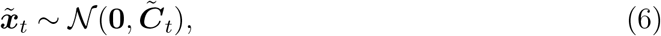

where 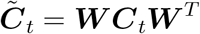. Due to the fact that ***W*** is full-rank and hence 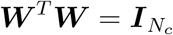,

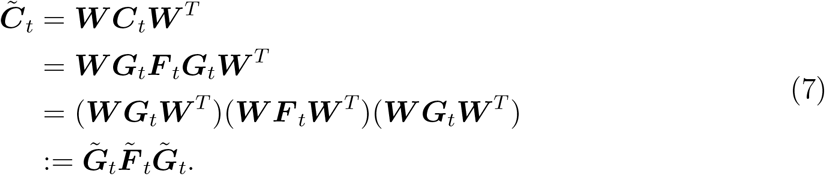

Then the projection matrix ***W*** can be passed in into linear combination equations:

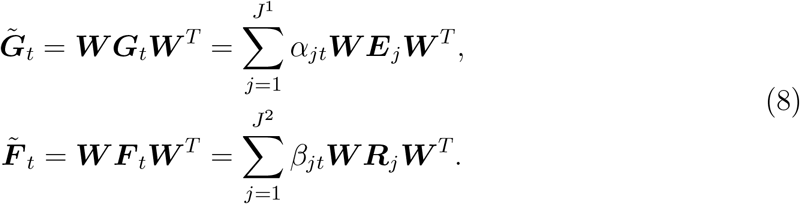

Now we have defined the generative model in the PCA-transformed space with the observation model parameters *θ*_*obs*_ in the original space.

### 6.5 Training M-DyNeMo

The goal of training is to infer the posterior distribution of the latent variables 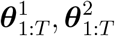 as well as the observation model parameters *θ*_*obs*_ = {***E***_*j*_ : *j* ∈ [*J* ^1^]} ∪ {***R***_*j*_ : *j* ∈ [*J* ^2^]}. This is achieved by minimising the variational free energy ([Blei et al., 2017]).

In practice, it is infeasible to feed the whole time series to the LSTM due to memory restrictions. More importantly, although the use of LSTM mitigates the issue of gradient explosion and vanishing to some extent, it is still an issue when the sequence is long. Hence we separate the data into *N*_*seq*_ sequences of length *T*_*seq*_ = 200 (so that ⌊*T/T*_*seq*_⌋ = *N*_*seq*_).

For M-DyNeMo, the loss for each sequence *n* is

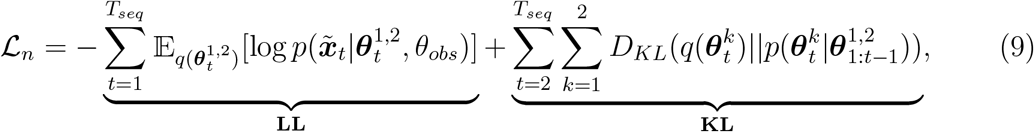

where *D*_*KL*_(·||·) is the Kullback–Leibler divergence ([Kullback and Leibler, 1951]) and we have used the mean field approximation:

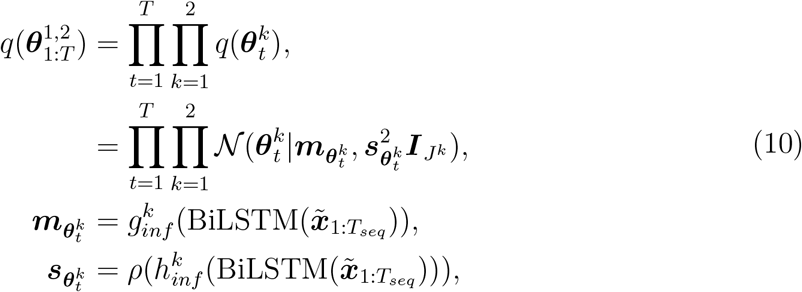

The learning of variational parameters are amortised by learning a map from the data to the variational parameters 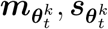 using bi-directional LSTM BiLSTM and learnable linear transformations 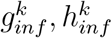. This allows not only scalable training, but also direct inference on unseen data. During training, mini-batches of sequences are shuffled and at each step of optimisation, each mini-batch of sequences is used to calculate the loss and gradient. Gradient descent based algorithm (in this paper ADAM, [Kingma and Ba, 2014]), together with the reparameterisation trick ([Kingma and Welling, 2013]) to pass gradient through the **LL** term, were used to minimise the loss.

Several tricks are used to help with the convergence of training and minimisation of the loss. See Section D for more details.

## A Simulation results

In this section, we describe how we use simulation to validate M-DyNeMo and investigate potential benefits of multi-dynamic modelling.

### A.1 Simulated data

We simulate data with the Hidden Markove Model (HMM) generative model. We prespecify the number of states *J* ^1^ = *J* ^2^ = *J*, the transition probability ***A*** and randomly simulate the observation model parameters *θ*_*obs*_, which are then used to simulate the observed data. Two simulation datasets are used:

- **Simulation 1**. Here we simulate data with multiple dynamics using *J* = 5, *N*_*c*_ = 40, *T* = 25600. The transition probability matrix is

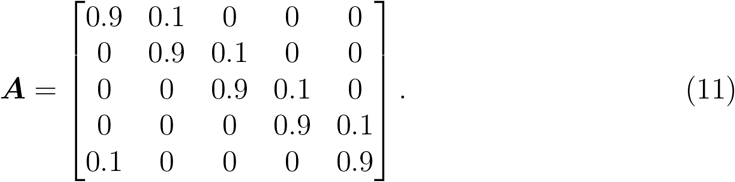
- **Simulation 2**. Here data is simulated with the same setting as Simulation 1 except that ***E***_*j*_, ***R***_*j*_ follow the same underlying temporal dynamics, i.e. they depend on the same underlying Markov chain of states.

### A.2 Simulation 1: M-DyNeMo performs more accurate inference on data with multiple dynamics

Here we aim to see if M-DyNeMo can perform inference more accurately than the traditional sliding window approach and DyNeMo, given the data has multiple dynamics. We focus on the first simulated data in Section A.1 and follow the procedure described in Section 6.3 with step size of 1 and window lengths ranging from 4 to 16. We show the DICE coefficients of the inferred time courses in Figure A.1a. We see the optimal window length of 6 samples does not give perfect DICE coefficient. If we look closely and compare the simulated and inferred time courses in Figure A.1b, we observe that sliding window approach does a decent job at inferring relatively long state changes, but fails to infer the transient state changes accurately. This is the reason why we turn to generative models that implicitly learn the time scale from data.

We now train M-DyNeMo and DyNeMo on the same simulated dataset. From Figure A.2a, we see that M-DyNeMo can recover both state time courses accurately while DyNeMo cannot, due to its intrinsic model assumption of shared dynamics. Specifically, DyNeMo only infers the correlation dynamics. A quantitative measure of the accuracy of inferred time courses measure by DICE coefficient is shown in Figure A.2b, where we see M-DyNeMo achieves perfect inference of both time courses. In this simulation study, the signal from time-varying power is relatively low compared to that from time-varying FC, which is why DyNeMo ignores the power dynamics. Additionally, we can investigate the reconstruction error measured by Riemannian distance between the reconstructed co-variances and the ground truth at each time point, shown in Figure A.2c. It is clear that M-DyNeMo has a lower reconstruction error than DyNeMo, indicating M-DyNeMo is a more accurate model of time-varying covariances on this data.

### A.3 Simulation 2: M-DyNeMo can infer accurately on single dynamics data

It is important to show that the separate dynamics is not an artefact invented by M-DyNeMo. This can be shown by training M-DyNeMo on the second simulated dataset where data were simulated with a single underlying dynamics shared by both standard deviation and correlation. Shown in Figure A.3a, both of the M-DyNeMo inferred mode time courses match the ground truth and the reconstruction error of time-varying covariances is comparable to that of DyNeMo, shown in Figure A.3b.

## B More results on real data

Here we show some additional results on real MEG data in Figures B.1, B.2, B.3, B.4, and B.5.

## C Generative model of M-DyNeMo

Here we give an illustration of the generative model of M-DyNeMo in Figure C.1.

## D Model training details

### D.1 Leanring correlation matrices

A square matrix ***R*** ∈ ℝ^*n×n*^ is a correlation matrix if and only if it is a covariance matrix with diagonal of 1. Similar to learning a covariance matrix, we cannot learn the entries of a correlation matrix directly and need to parameterise the matrix with a vector using a bijection. Similar to learning a covariance matrix, we resort to learning the Cholesky decomposition of a correlation ***R*** = ***LL***^*T*^, where the *k, l* element ***L***_*kl*_ satisfies:

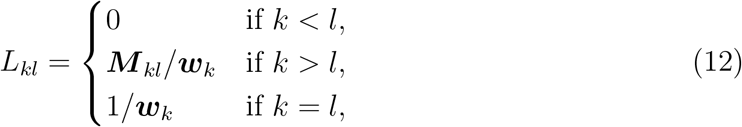

where ***M*** _*kl*_ is the *k, l* entry of a lower triangular matrix ***M*** and ***w*** is a vector such that its *k*-th element is 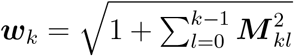. It is easy to check that ***R*** constructed in this way is a legitimate correlation matrix and during training, we learn the vector of lower triangular entries of ***M***.

**Figure A.1:**
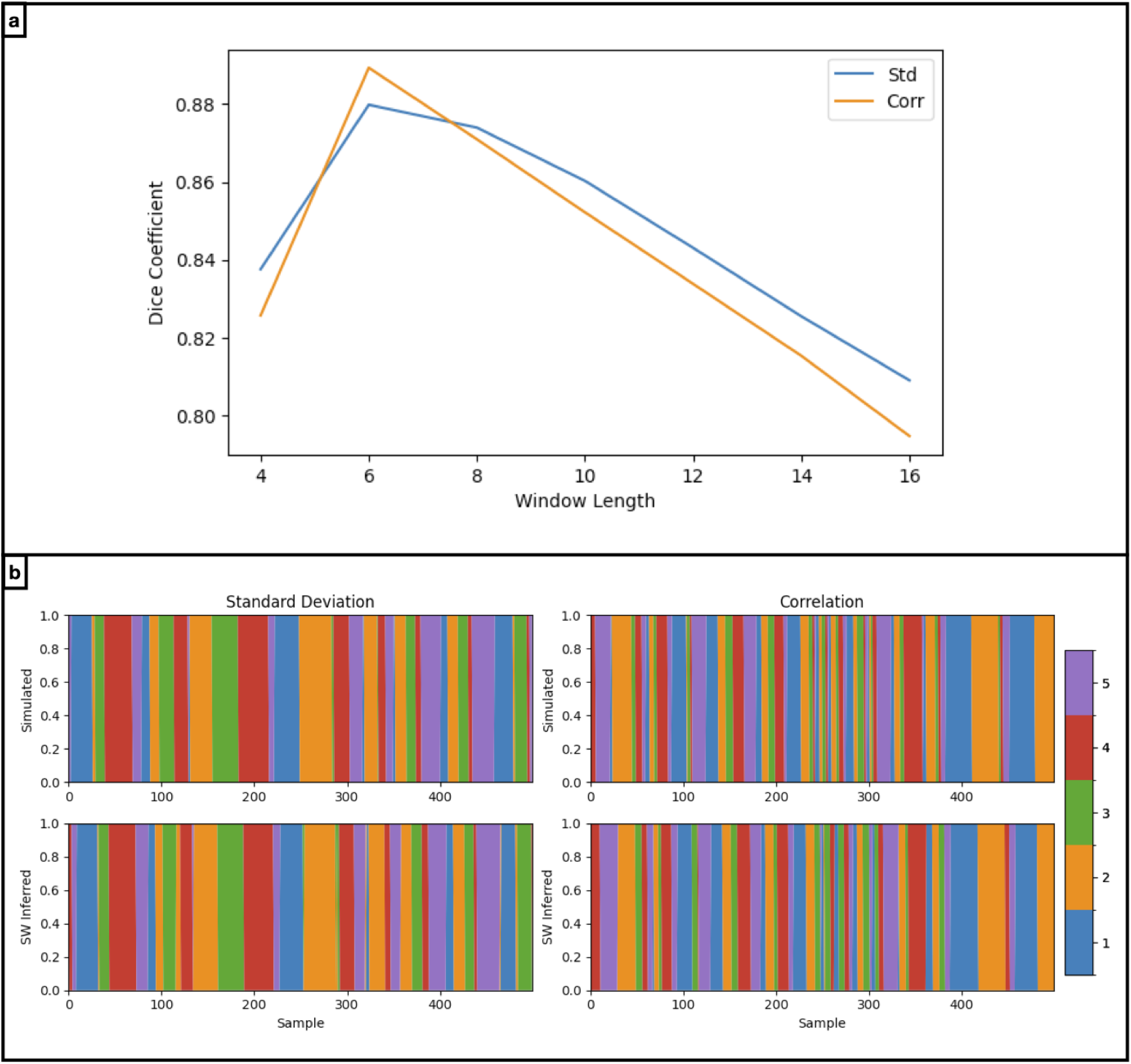
Sliding window approach fails to infer ground truth state time courses. *Results on Simulation 1 dataset*. a) Dice coefficient of state time courses inferred by sliding window approach using different window lengths. b) The simulated (top row) and sliding window approach inferred (bottom row, with window size of 6 samples) time courses. A segment of 500 samples containing short state transitions is shown.

**Figure A.2:**
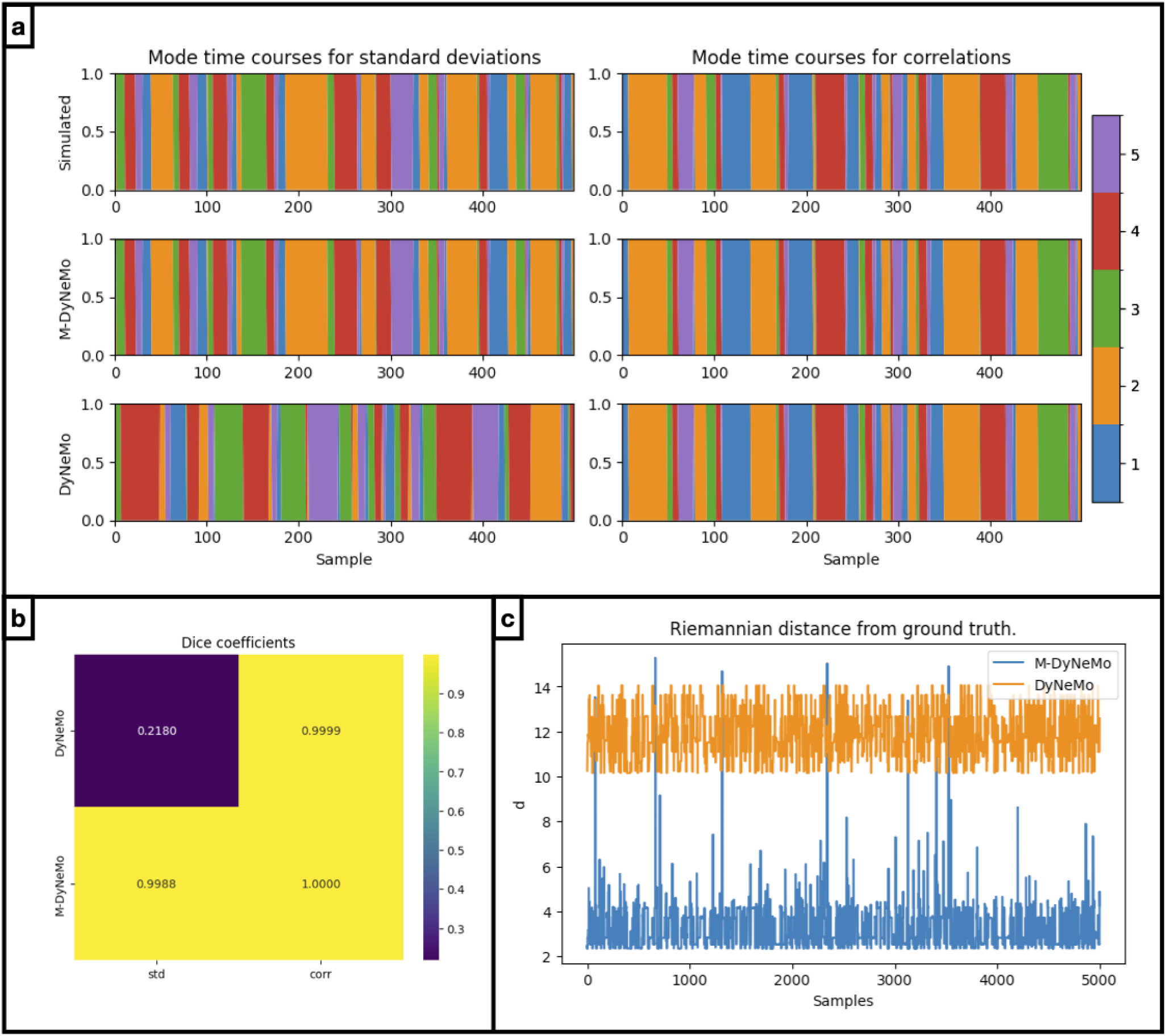
M-DyNeMo infers time courses more accurately than DyNeMo when data has multiple dynamics. *Results on Simulation 1 dataset*. a) The simulated (top row), M-DyNeMo inferred (middle row) and DyNeMo inferred (bottom row) time courses for time-varying standard deviations (left column) and correlations (right column). Here different colours indicate activation of different states and only the first 500 samples are plotted. b) DICE coefficients of DyNeMo and M-DyNeMo inferred time courses compared to the simulated ground truth. c) The reconstruction error (Riemannian distance) of the first 500 samples of time varying covariances is plotted against sample on the x-axis. The reconstruction error of DyNeMo is plotted in orange and that of M-DyNeMo is plotted in blue.

**Figure A.3:**
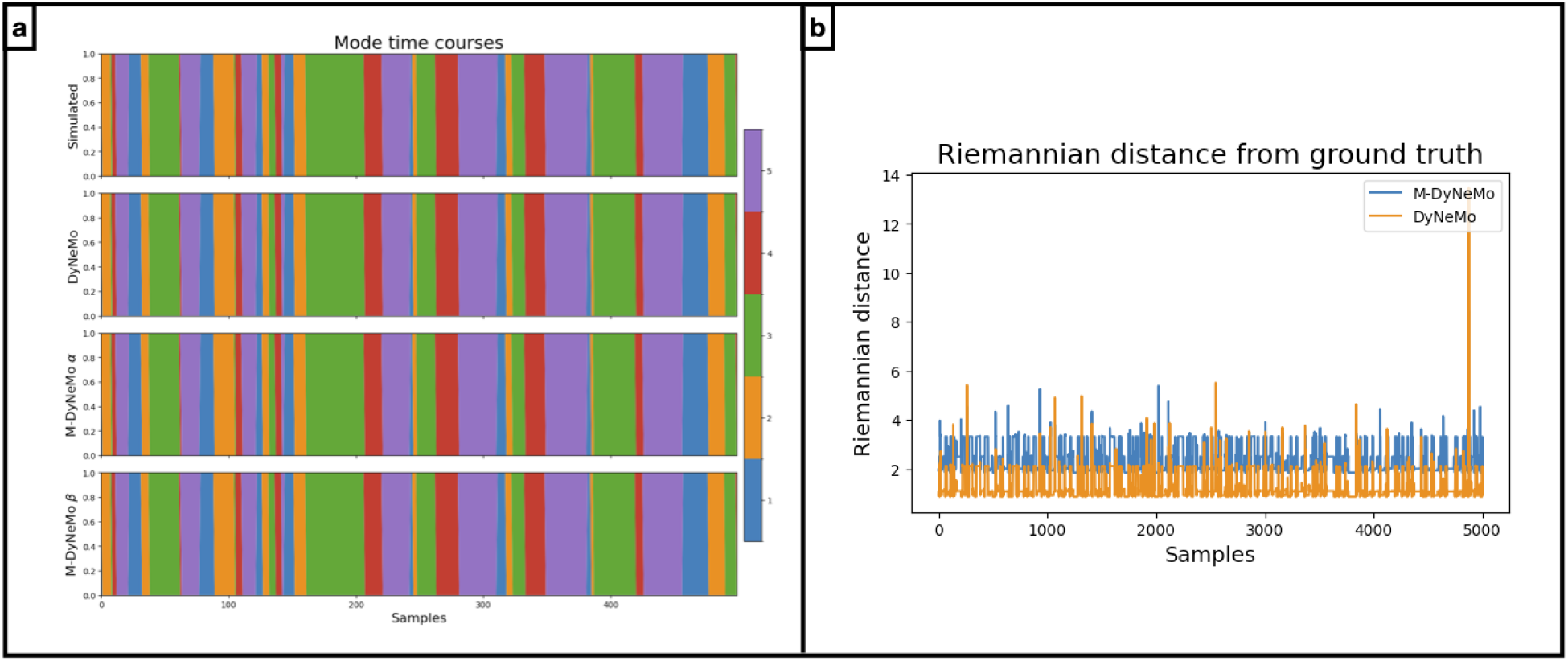
M-DyNeMo infers accurately on data with single dynamic. *Results on Simulation 2 dataset*. a) From top to bottom row are the ground truth simulated, the DyNeMo inferred, and the two “separate” M-DyNeMo inferred mode time courses. Here different colours indicate activation of different states and only the first 500 samples are plotted. b) The reconstruction error (Riemannian distance) of the first 5000 samples of time-varying covariances is plotted against sample on the x-axis. The reconstruction error of DyNeMo is plotted in orange and that of M-DyNeMo is plotted in blue.

**Figure B.1:**
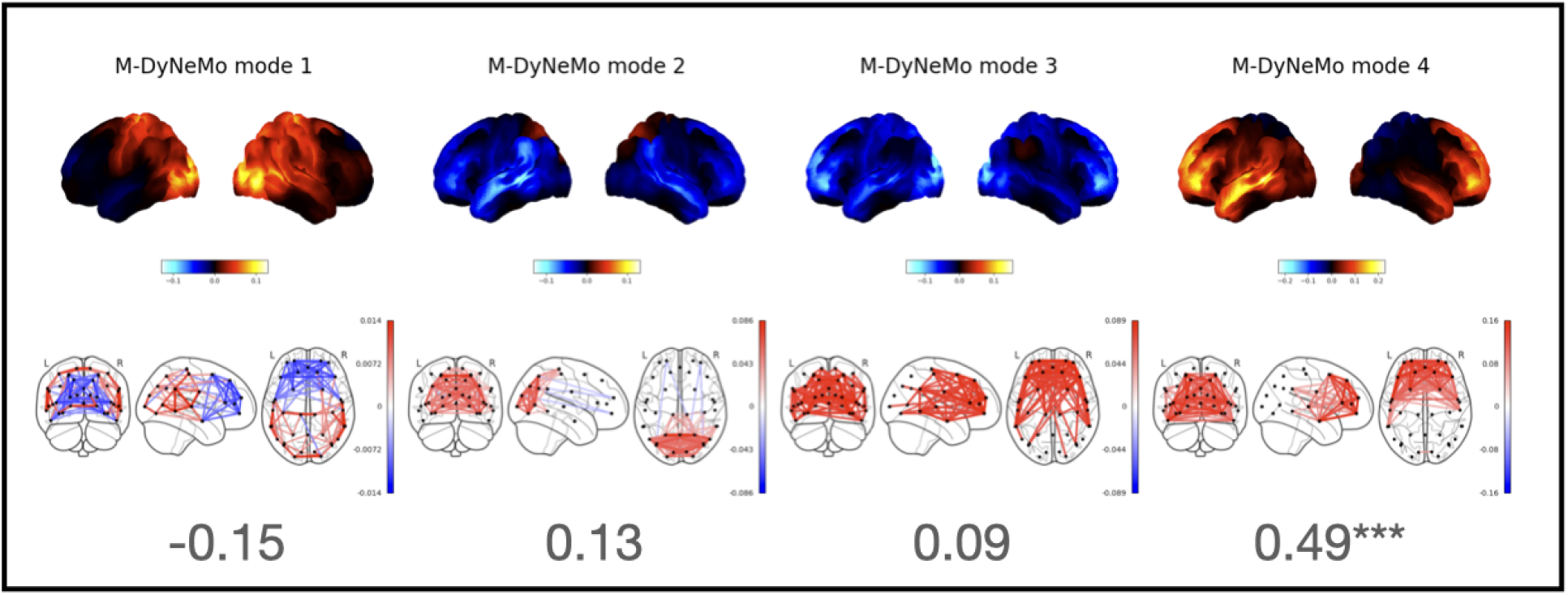
Single dynamic DyNeMo is dominated by time-varying power. *Results from amplitude time courses (1-45Hz) of 38 brain regions obtained from MEGUK-38 resting-state data*. Power (top row) and FC (covariance, bottom row) network maps given by re-calculated networks using M-DyNeMo inferred FC mode time course *β*. The cosine similarity is also shown for each pair of modes between the re-calculated covariances on M-DyNeMo inferred FC mode time course *β* and standard DyNeMo inferred covariances.

**Figure B.2:**
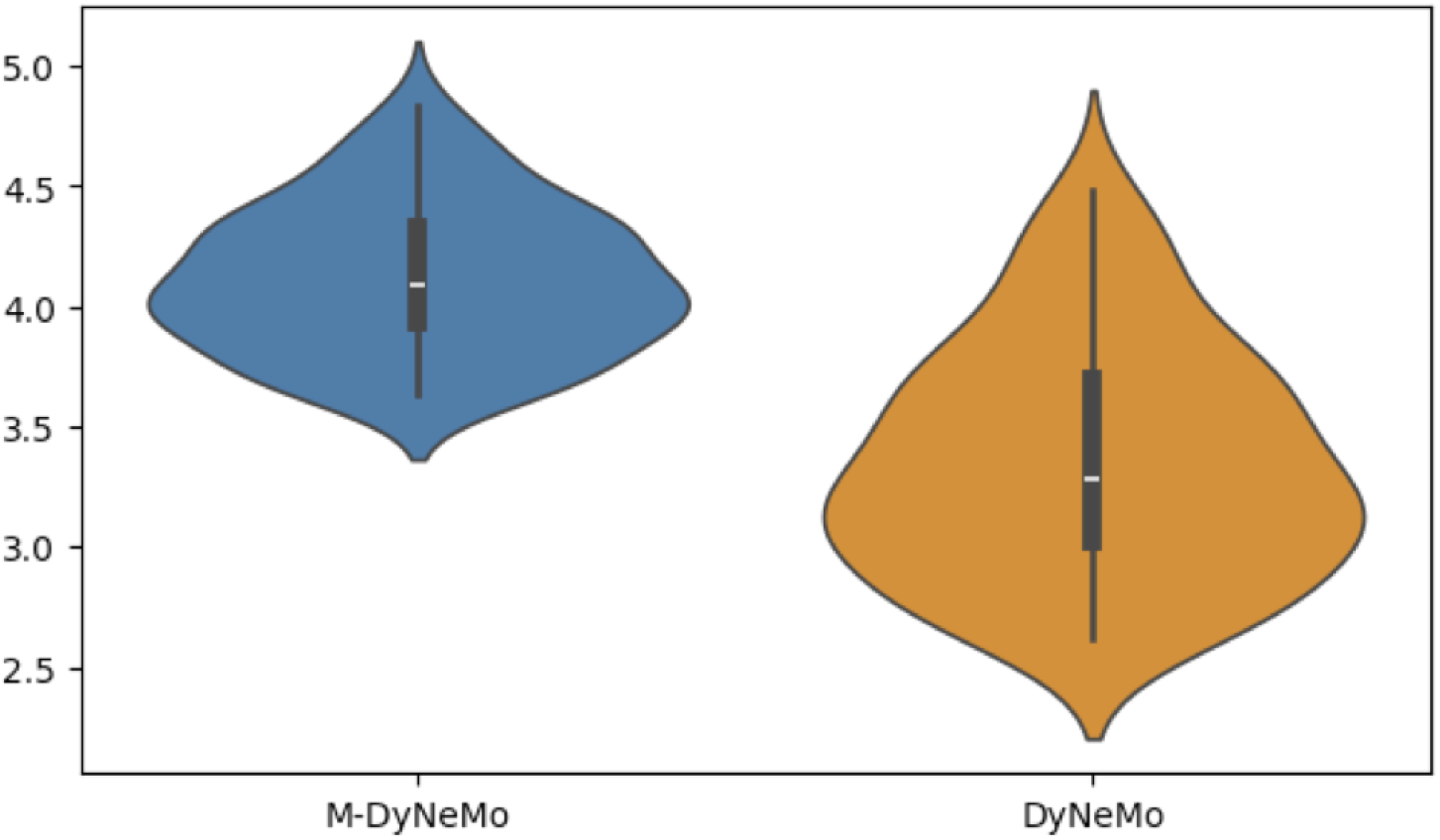
M-DyNeMo reconstructed time varying covariances show greater fluctuation over time. *Results from amplitude time courses (1-45Hz) of 38 brain regions obtained from MEGUK-38 resting-state data*. For M-DyNeMo and DyNeMo, the y-axis shows the variability of model reconstructed time varying covariances in terms of Riemannian distance of time varying covariances from the mean covariance for each subject.

**Figure B.3:**
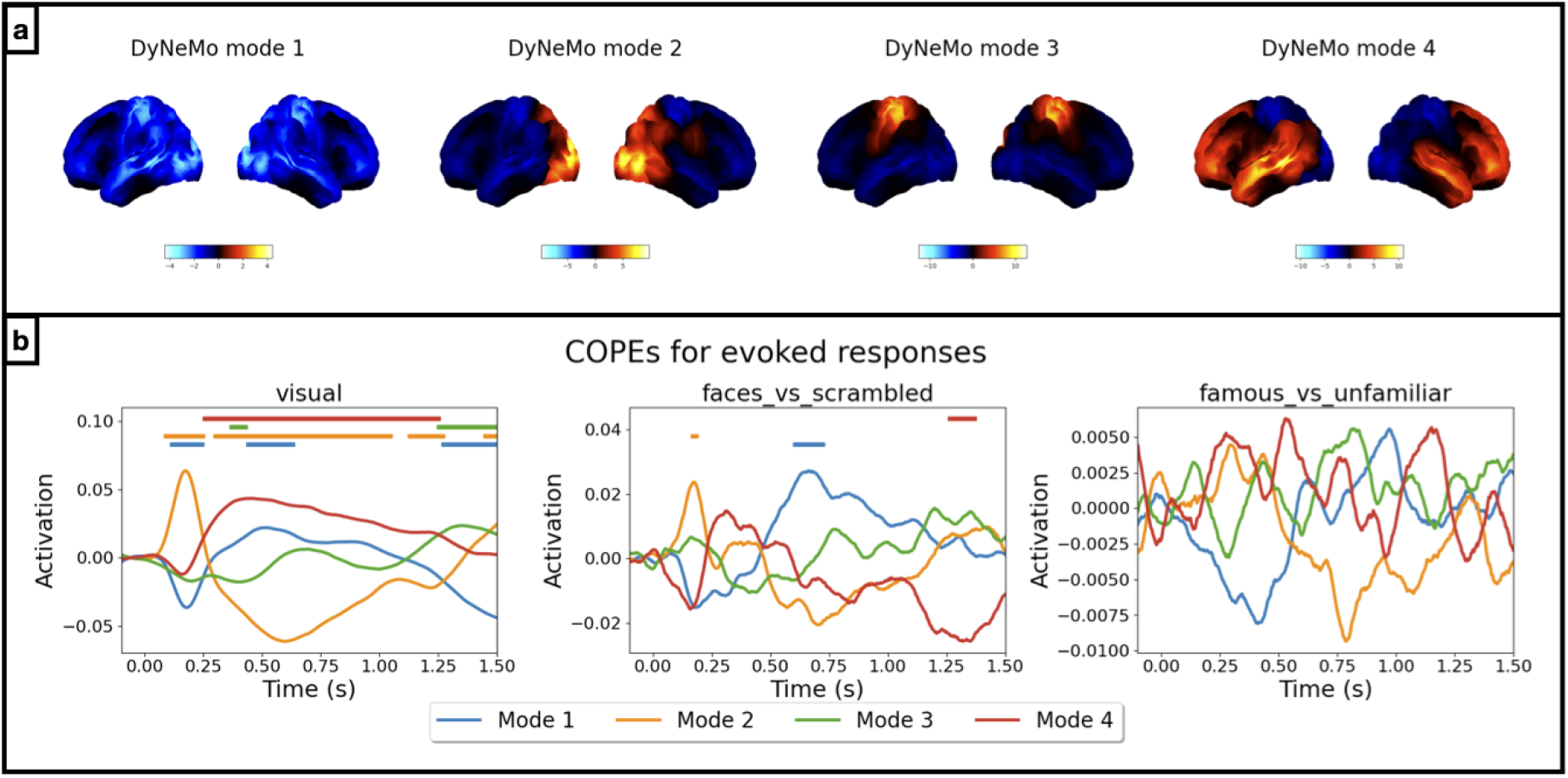
Evoked responses given by DyNeMo. *Results from amplitude time courses (1-45Hz) of 38 brain regions obtained from Wakeman-Henson data*. a) Power maps for different modes given by DyNeMo. b) Evoked response analysis on DyNeMo-inferred mode time course. Different colours show baseline corrected evoked response (contribution to overall variance) of different modes.

**Figure B.4:**
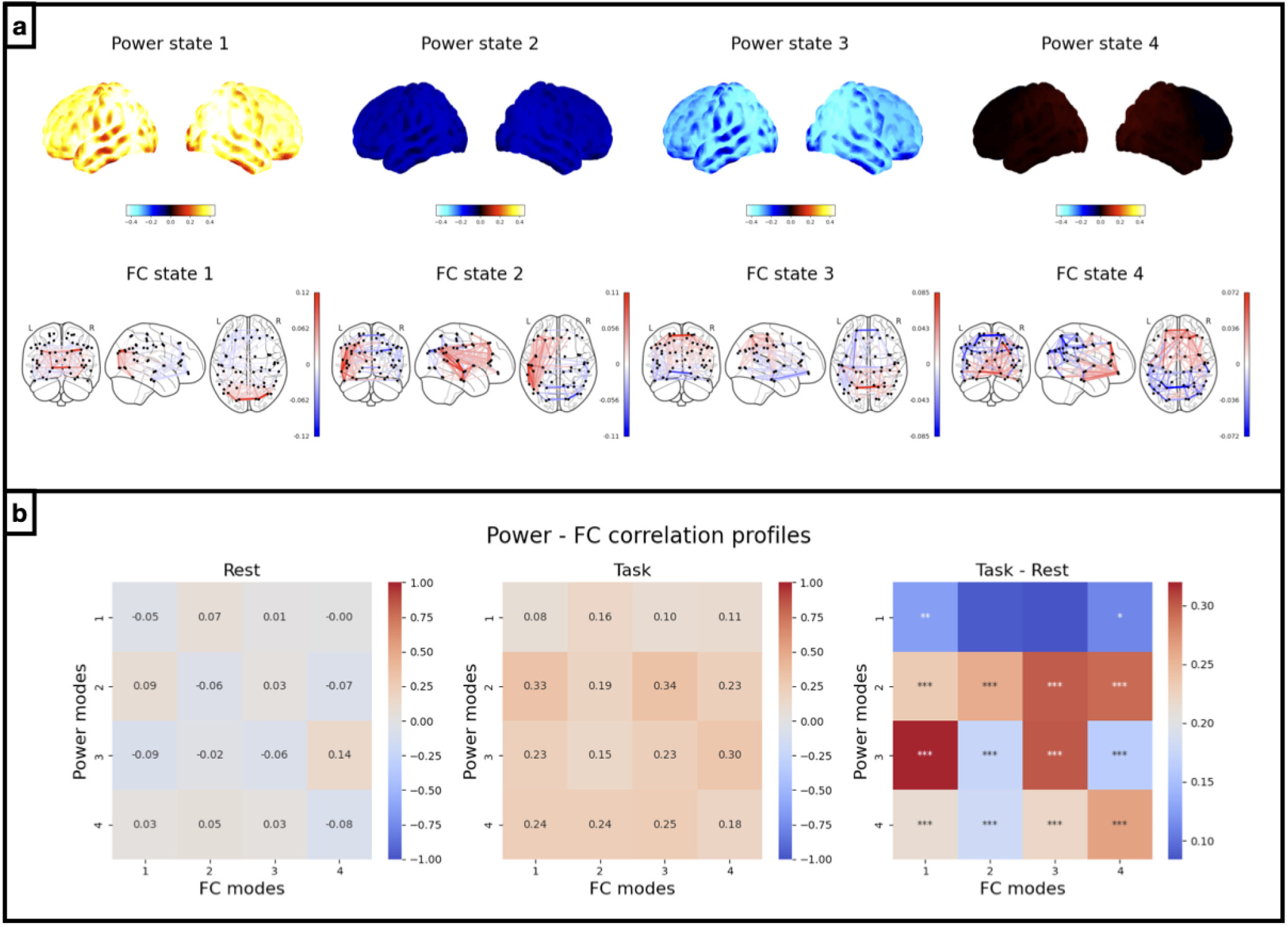
Sliding window approach reveals evidence of different correlation profiles during task versus during rest. *Results from amplitude time courses (1-45Hz) of 52 brain regions obtained from MEGUK-52 data*. a) Power (top row) and FC (bottom row) maps for each mode of the two state time courses. b) Correlations within *α* (left) and within *β* (middle) are shown. The right plot shows the differences in correlations during task versus during rest. Correlations that significantly differ from zero are marked with asterisks depending on the p-values.

**Figure B.5:**
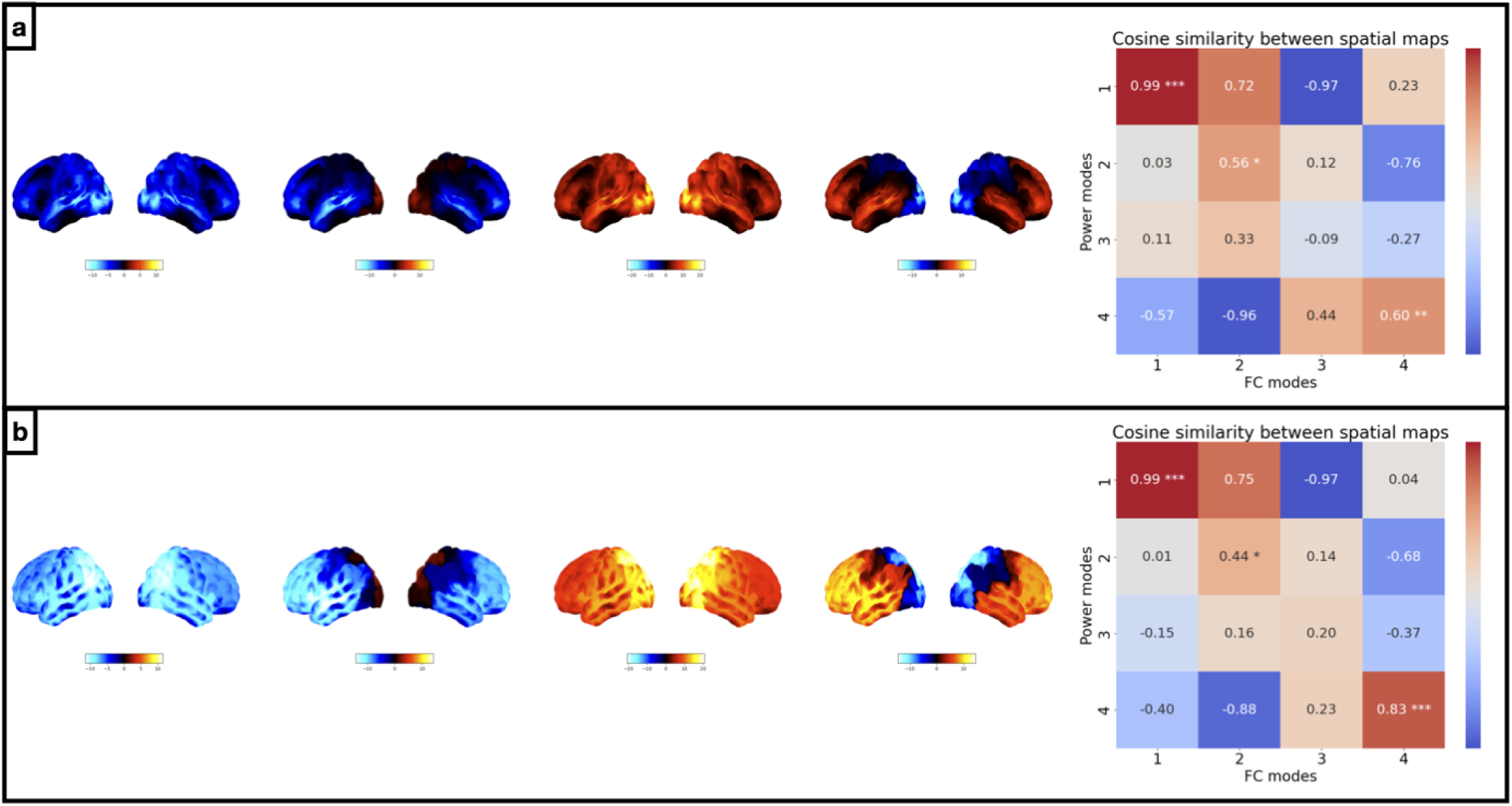
Additional FC degree spatial maps. *Results from amplitude time courses (1-45Hz) of 38 brain regions obtained from Wakeman-Henson data and of 52 brain regions obtained from MEGUK-52 data*. a) FC degree spatial maps on the Wakeman-Henson dataset (left) and the cosine similarity between power maps and FC degree maps for each pair of power and FC modes (right). b) FC degree spatial maps on the MEGUK-52 dataset (left) and the cosine similarity between power maps and FC degree maps for each pair of power and FC modes (right).

**Figure C.1:**
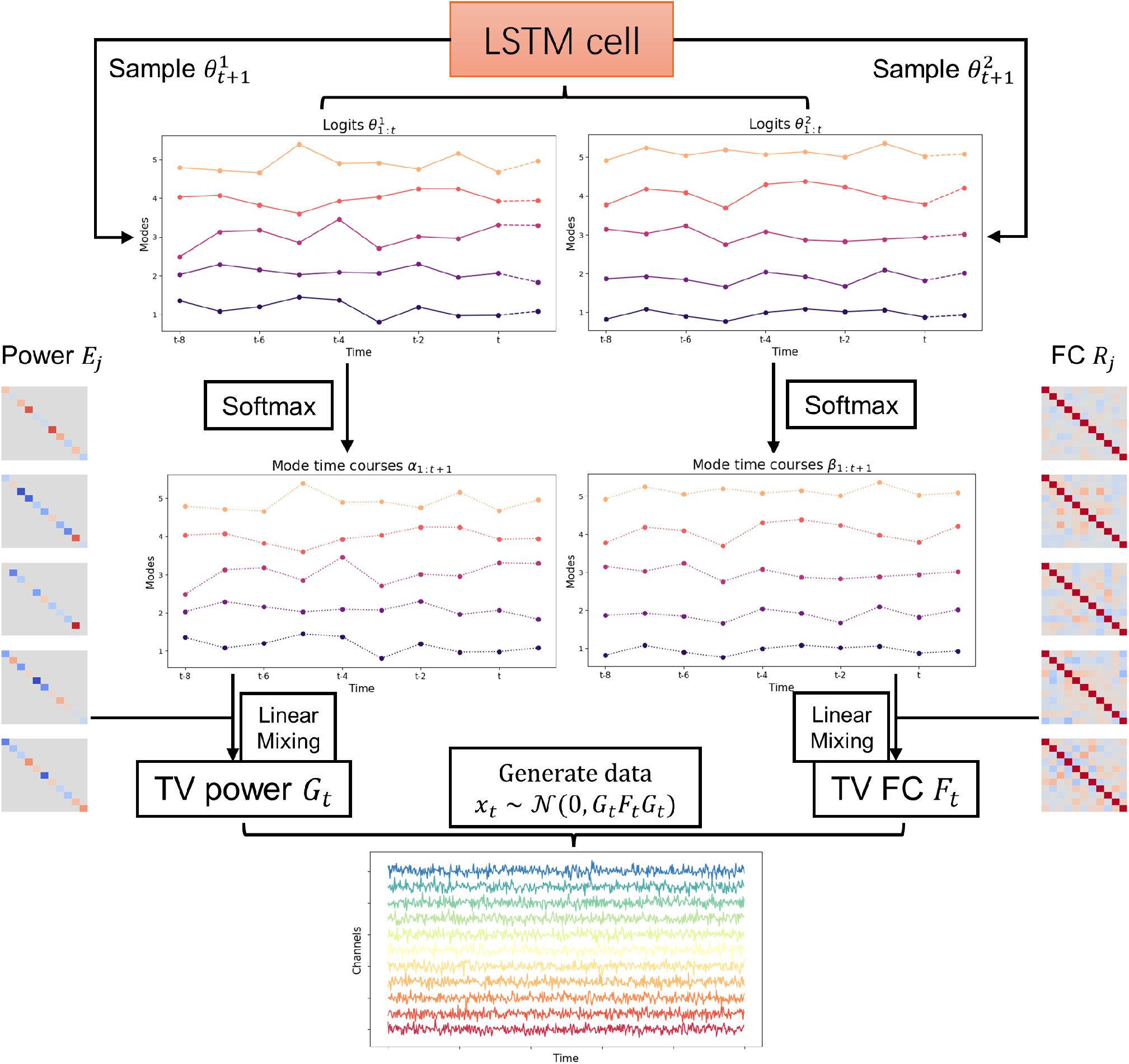
Generative model of M-DyNeMo. Historic logits 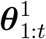 and 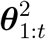 are concatenated and passed to the LSTM cell. The outputs of the LSTM cell are the parameters of the distribution from which we sample the future logits 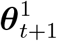 and 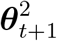. The sampled logits are concatenated to the original logit sequences and the process is repeated. The generated sequences of logits are normalised with a softmax transformation and we get the mode time courses ***α***_1:*T*_, ***β***_1:*T*_. Diagonal matrices ***E***_*j*_ and correlation matrices ***R***_*j*_ are linearly mixed with ***α***_1:*T*_ and ***β***_1:*T*_ respectively to give the time-varying power ***G***_*t*_ and FC ***F***_*t*_. Finally the observed ata are generated through the multivariate Gaussian distribution with time-varying covariance ***G***_*t*_***F***_*t*_***G***_*t*_.

### D.2 Extra tricks for model training

Some extra tricks were used to help fast and stable convergence of M-DyNeMo:

- **KL annealing**: M-DyNeMo can be thought of as a special case of the variational autoencoder (VAE) and optimising a VAE with RNN is challenging. In most runs, the **KL** term dominates and the model consistently sets the variational distribution close to the prior distribution of the latent variables, yeilding very small **KL** term ([Bowman et al., 2015]). The soluation is surprisingly simple and effective: annealing the **KL** term. The idea is to train the model with a modified loss function where the **KL** term is multiplied by an annealing factor *κ* that starts at 0 and gradually increases to 1 as the training progresses, i.e.

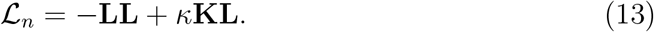 This allows the model to first learn a good encoder before tuning the variational distribution to match the prior distribution.
- **Multi-start initialisation**: The objective function is highly non-convex and the model can end up in different local minima depending on the initialisation. To mitigate this, a multi-start initialisation strategy is used. At each initialisation, the model parameters are initialised randomly and the model is trained for a small number of epochs. This is repeated multiple times and the model with the lowest loss is chosen for further training.
- **Learning rate scheduling**: A learning rate scheduler is used to exponentially reduce the learning rate as the training progresses. This helps the model to take large steps in the beginning for faster convergence and smaller steps towards the end for fine-tuning. In practice, we only start decreasing the learning rate after the KL annealing phase.
- **Initialisation for *R***_*j*_: We find in practice that without a “good” initialisation of the correlation matrices, the convergence of the model is extremely slow. To help with this, we initialise the correlation matrices with “realistic” ones obtained from the data. Specifically, we use a sliding window approach to calculate the time-varying correlation matrices and used K-means clustering to identify *J* ^2^ clusters of time-varying correlation matrices. We then initialise the correlation matrices with the cluster centroids. We find that this helps the model to converge to a lower loss consistently.

## E Post-hoc analysis

### E.1 Re-normalising mode time courses

As mentioned in [Gohil et al., 2022], the inferred mode time courses do not account for the difference in the relative magnitude of each of the spatial modes. For *α*, the raw mode time course is multiplied by the mean of the square of the diagonal elements of ***E***_*j*_ before normalised by dividing the sum over modes at each time point. Similarly for *β*, the raw mode time course is multipled by the mean of absolute off-diagonal elements of ***R***_*j*_ and normalised.

### E.2 Maximum statistic permutation test for spatial map similarity

The underlying assumption of this test is that sequences of mode time courses are exchangeable under the null hypothesis and we used the maximum statistic approach to correct for multiple comparison. The procedure for generating samples from the null distribution is as follows:

- Separate mode time courses into sequences of 1 second.
- Shuffle the sequences.
- Regress TV variances/correlations/covariances on the shuffled mode time courses and get regressed spatial maps.
- Match the spatial maps with cosine similarity using the Hungarian algorithm ([Kuhn, 1955]).
- Get the maximum cosine similarity across pairs of modes.

The p-value of the cosine similarities of each pair of modes is calculated as the percentage of samples from the null distribution that exceeds the observed cosine similarity. In this paper, tests with p-value lower than 0.05, 0.01, and 0.001 are annotated with one, two, and three asterisks respectively.

